# Replication origins drive genetic and phenotypic variation in humans

**DOI:** 10.1101/2021.12.16.472898

**Authors:** Pierre Murat, Guillaume Guilbaud, Julian E. Sale

## Abstract

DNA replication starts with the activation of the replicative helicases, polymerases and associated factors at thousands of origins per S-phase ^1^. Due to local torsional constraints generated during licensing ^2^ and the switch between polymerases of distinct fidelity and proofreading ability following firing ^3,4^, origin activation has the potential to induce DNA damage and mutagenesis. However, whether sites of replication initiation exhibit a specific mutational footprint has not yet been established. Here we demonstrate that mutagenesis is increased at early and highly efficient origins. The elevated mutation rate observed at these sites is caused by two distinct mutational processes consistent with formation of DNA breaks at the origin itself and local error-prone DNA synthesis in the immediate vicinity of the origin. We demonstrate that these replication-dependent mutational processes create the skew in base composition observed at human replication origins. Further, we show that mutagenesis associated with replication initiation exerts an influence on phenotypic diversity in human populations disproportionate to the origins’ genomic footprint: by increasing mutational loads at gene promoters and splice junctions the presence of an origin influences both gene expression and mRNA isoform usage. These findings have important implications for our understanding of the mutational processes that sculpt the human genome.

## MAIN

Mutations are the building blocks of genome evolution and the driving force for both Mendelian disorders and cancer. Detailed catalogues of human genetic variation in normal cells and tumours have begun to reveal some of the determinants of mutation distribution in human cells. Large-scale genomic features, including replication timing ^5^ and chromatin structure ^6^ influence the rate of germline and somatic mutations at the megabase scale. Mutation rates are also influenced by local features such as nucleosome positioning ^7^, the orientation of the DNA minor groove around nucleosomes ^8^, transcription factor binding ^9,10^ and transcription initiation ^11^. These analyses, and others ^12^, have established a principle that differential DNA accessibility, due to DNA condensation or active biological processes, significantly contributes to variable local mutation rates in the human genome. DNA replication initiation involves several events with the potential for local DNA damage and mutagenesis, including activation of the replicative helicases and origin DNA melting ^13^, and a series of polymerase switches ^4^. This suggests that origins may be hotspots for mutagenesis, and that local mutation rates should reflect the usage and efficiency of individual origins. However, the lack of high-resolution methods for mapping replication origins and quantifying initiation efficiency have hampered the characterisation of the nature and consequences of the mutational processes operating at replication origins.

### Origin mutation rate analysis

To determine the impact of replication initiation on mutation rates, we took advantage of recent high-resolution maps of human replication origins. We first categorised human replication origins according to their usage and efficiency. A map of origins, obtained with a Short Nascent Strand isolation protocol coupled with next-generation sequencing (SNS-seq), from 19 human cell samples ^14^ representing untransformed and immortalised cell types allowed determination of origin usage. This approach identified 320,748 origins of which 256,600 were found to be cell type dependent and 64,148 were found in all analysed cell lines, hereafter referred to as stochastic and core origins respectively (**Fig. 1a**). We refined this dataset by annotating origins enriched in early replicating domains of the human genome identified with our own high-resolution origin mapping technique, ini-seq 2 ^15^. Ini-seq 2 is a modification of initiation site sequencing (ini-seq) ^16^ that as well as providing high-resolution origin position information, gives a quantitative estimate of initiation efficiency, *i*.*e*. the probability that an identified origin fires at each cell division. Ini-seq 2 identified 23,905 core origins, at a resolution below 1 kb, which are enriched in conserved early replication domains of the human genome (**Extended Data Fig. 1a**), and that are hereafter referred to as constitutive origins. Importantly, as our ini-seq 2 method allows the categorisation of human replication origins by efficiency, we could demonstrate that origin base composition determines origin usage and efficiency (**Extended Data Fig. 1b, c**).

**Figure 1.**
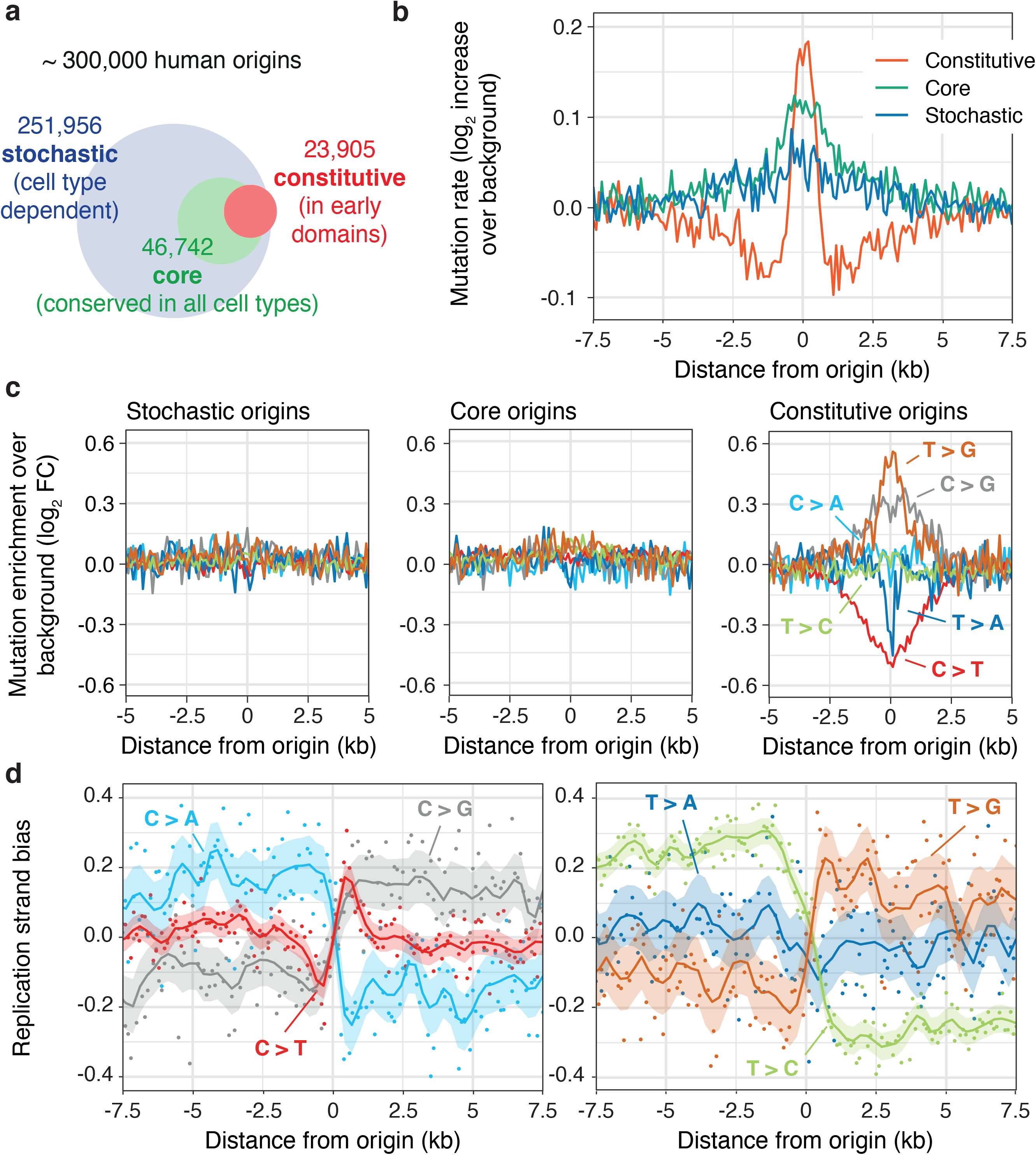
Elevated mutation rates at constitutive origins. **a**, Categorisation of human replication origins based on usage. Stochastic origins ^14^ are cell type specific, while core ^14^ and constitutive ^15^ origins are replication initiation hotspots independent of cell type. Constitutive origins are enriched in early replicating domain of the genome. **b**, Mutation rates associated with population SNPs are increased at all origin types compared to their flanking domains (*P* = 2.46 × 10^−4^, 3.50 × 10^−8^ and 0.014 for constitutive, core and stochastic origins respectively, chi-square test). Mutation rates are reported as the increase over background values computed from domains adjacent to origins. **c**, Increased mutation rates at stochastic and core, but not constitutive origins are a consequence of local sequence composition effects. Correcting mutation rates by the local base composition show an excess of T > G, C > G, T > C and to some extent C > A mutations at constitutive origins. The mutated base is represented by the pyrimidine of the base pair. **d**, Mutagenesis at constitutive origins is replication dependent as the excess mutations at constitutive origins exhibit a replication strand bias. Replication strand bias is computed as the ratio between the density of mutation at a given base pair over the density of mutation at the complementary base pair.

We then quantified mutation rates by assessing the distribution of common single nucleotide polymorphisms (SNPs) and small insertions/deletions (indels) at replication origins. We found increased SNP frequency and decreased distances in between SNPs at all origin types compared with their flanking regions (**Fig. 1b** and **Extended Data Fig. 1d**). Mutation frequency for long (≥ 2 bp) but not short (= 1bp) indels was increased at core and constitutive origins (**Extended Data Fig. 1e**). However, when correcting for local variation in base composition we found that the apparently elevated SNP mutation rate observed at stochastic and core origins is explained by the sequence context (**Fig. 1c**). For constitutive origins though, the observed excess of T > G, C > G, T > C and to some extent C > A mutations was not accounted for by local changes in base composition (**Fig. 1c**). It is notable that C > T mutations, the most frequent mutation in human populations ^17^, are depleted at constitutive origins. The four enriched mutation types all exhibited significant strand biases (**Fig. 1d**), the polarity of which inverts at the origins, supporting a role for DNA replication in the generation of these mutations. Further, the magnitude of the strand bias was dependent on the efficiency of the constitutive origins (**Extended Data Fig. 1f**). Such replication strand bias was not observed at stochastic and core origins (**Extended Data Fig. 1g**). These observations suggest a specific form of replication-dependent mutagenesis is focussed on constitutive origins.

### Replication origin mutation signatures

To characterise these mutational processes, we identified the mutational signatures operating at constitutive origins using an established non-negative matrix factorization (NMF) approach ^18^. We first extracted single-base substitution signatures (SBS) from SNP mutation count matrices reporting the observed frequencies of the 6 pyrimidine base substitutions in their trinucleotide context. We identified three signatures, which we termed SBS A, B and C, operating in the vicinity of constitutive origins (**Fig. 2a**) and analysed their relative contribution to mutagenesis at these origins (**Fig. 2b**). While SBS A mainly operates outside constitutive origins, SBS B and C contribute to mutagenesis within a ~ 4 kb domain around them. SBS B operates in a narrow 1 kb window centred on the origins, while SBS C displays a ‘volcano-shaped’ profile extending 2 kb either side of the origins with a strong depletion exactly at their centre. The contribution of SBS B and C to mutagenesis at origins correlates with constitutive origin efficiency (**Extended Data Fig. 2a, b**) and reflects origin usage (**Extended Data Fig. 2c**) suggesting that they are promoted by replication initiation.

**Figure 2.**
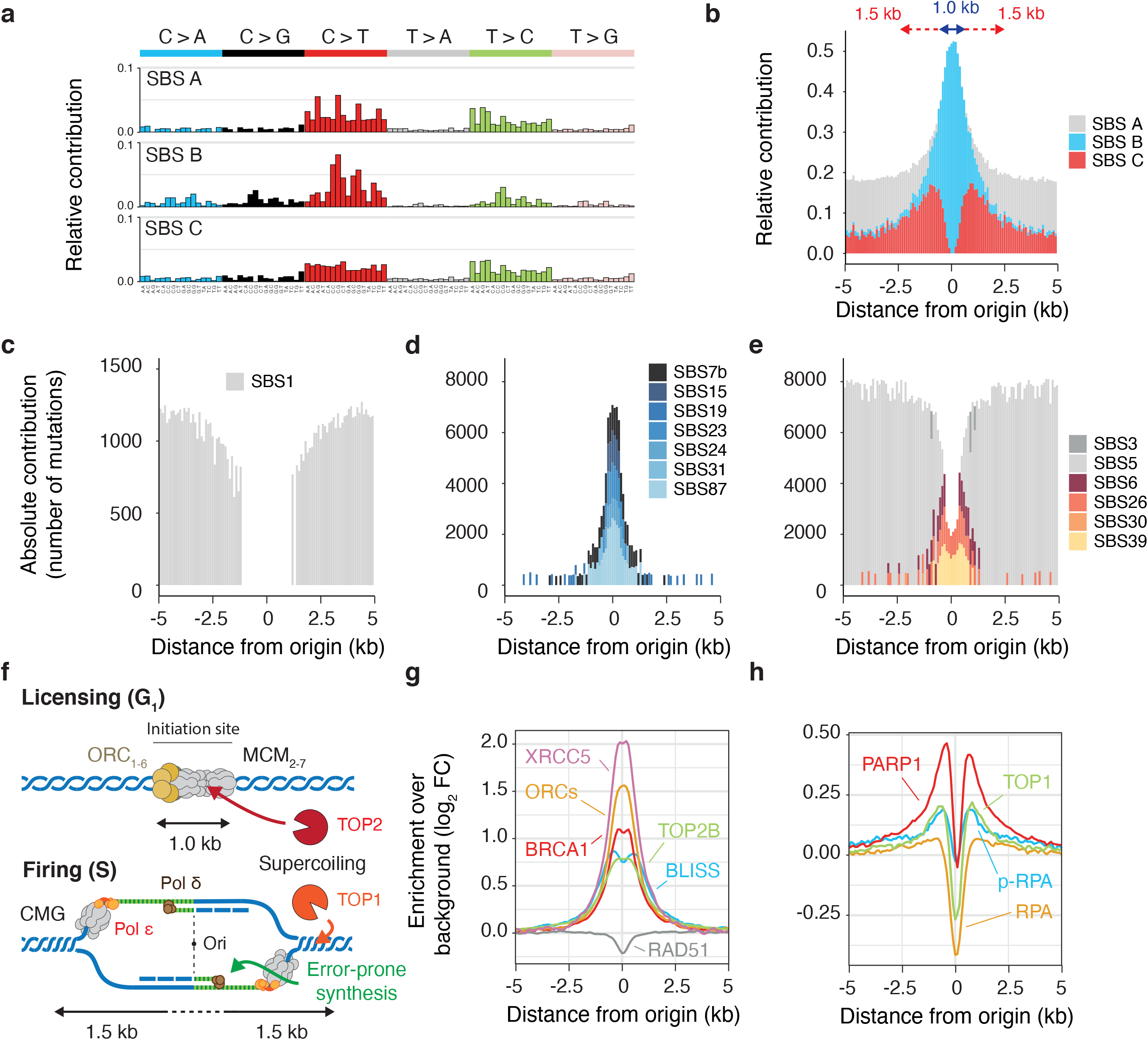
Distinct mutational processes operate at constitutive origins. **a**, An unbiased *de novo* mutational signature analysis identifies three single-base substitution signatures, SBS A-C, describing mutation patterns at constitutive origins. Each profile shows the mutation probability of each indicated context-dependent base substitution type. **b**, Assessing the relative contribution of SBS A-C in the vicinity of constitutive origins shows that, while SBS A mainly operates outside origins, SBS B and C contribute to mutagenesis within a ~ 4 kb domain centred at the origins. Signature exposures are reported as the relative contribution of each signature in 100 bp windows around constitutive origins. **c-e**, To characterise the mutational processes operating at constitutive origins, SBS A-C were compared to known signatures of somatic mutations in noncancerous and cancerous human tissues. **c**, The origin-associated signature SBS A is mainly explained by the known signature SBS 1, a clock-like background mutational process. **d**, SBS B is reconstructed from a set of 7 known signatures associated with DNA damage. **e**, SBS C is reconstructed from a set of 6 known signatures including SBS 5, describing another clock-like background mutational process, and signatures linked with defective MMR. **f**, A model for mutagenesis at constitutive origins depicting putative sources for mutagenesis during origin licensing (G1-phase) and firing (S-phase). Key steps of the model were validated by probing for the presence of DNA damage and repair factors using available ChIP-seq or genomic datasets, at the centre (**g**) or the flanks (**h**) of constitutive origins. Reported signals are averages from experiments conducted in different cell types when available. Enrichments were computed in 100 bp windows and normalised using background values from domains adjacent to origins.

We next identified three small insertion/deletion (indel) signatures, hereafter referred to ID A, B and C, operating in concert with SBS A, B and C respectively (**Extended Data Fig. 2d, e**). We found that while the ID A and C signatures are mainly composed of insertions and deletions at long (≥ 5 bp) homopolymers, indicative of DNA polymerase slippage and incomplete DNA mismatch repair (MMR) ^19^. In contrast, the ID B signature is characterised by increased long (≥ 5 bp) deletions with at least 5 bp of microhomology at their boundaries, indicative of DNA double-strand breaks (DSBs) processed by alternative end-joining (alt-EJ) mechanisms ^20^, or microhomology-mediated break-induced replication (MMBIR) ^21^.

To shed light on the molecular mechanisms underlying mutagenesis at constitutive origins, we compared the extracted SBS A-C signatures to known signatures of somatic mutations in noncancerous and cancerous human tissues ^22^. We identified a set of 14 known signatures that most closely reconstruct the mutation profiles of SBS A-C (**Extended Data Fig. 3a**) and assigned each of these signatures to one of the origin-associated signatures based on their cosine similarity and exposure. Our SBS A signature is mainly explained by the known SBS 1 signature (**Fig. 2c**), caused by the spontaneous or enzymatic deamination of 5-methylcytosine (5mC), and described as a clock-like mutational process that leads to the accumulation of mutations in all cell types in a time-dependent manner ^23^. Hence SBS A can be considered as a background mutational process. A set of 7 signatures, comprising SBS 7b (UV exposure), SBS 24 (aflatoxin exposure) and SBS 31 (platinum chemotherapy treatment) allows the reconstruction of the SBS B profile operating at constitutive origins (**Fig. 2d**). This set of signatures describes mutagenic processes associated with bulky DNA damage that can lead to replication fork stalling, collapse and DNA breaks ^24^. Finally, a set of 6 SBS signatures reconstruct the mutagenic processes operating in the flanks of constitutive origins, displaying the ‘volcano-shaped’ profile of SBS C (**Fig. 2e**). This set of signatures includes SBS 5, another background clock-like process ^23^, and SBS 6 and SBS 26, two signatures associated with defective MMR ^22^. Exposure of origins to the mutational processes defined by these SBS signatures is replication-dependent as their contribution to origin mutagenesis is usage and efficiency-dependent (**Extended Data Fig. 3b** and **3c**). Taken together these observations suggests that SBS B and C are likely to originate from unrepaired DNA breaks and persistent mismatches induced by replication initiation.

### A model for mutagenesis at origins

These results lead us to propose a model, summarised in **Fig. 2f**, to explain the mutagenesis observed at constitutive origins. We validated key features of our model by analysing available ChIP-seq datasets phasing the signals for key proteins on constitutive origins (**Fig. 2g** and **2h**) and assessing the impact of origin efficiency on protein enrichment around those origins (**Extended Data Fig. 3e**).

During the G1 phase of the cell cycle, licensing of all potential replication origins starts with the binding of the origin recognition complex (ORC) which displays high affinity for negatively supercoiled DNA ^25^ and binds to origins along with DNA topoisomerase II ^26,27^. Consistent with this, we found that ORC proteins colocalise with TOP2B at constitutive origins to define ~ 1 kb replication initiation sites (**Fig. 2g**). Abortive topoisomerase II activity at origins leads to potentially mutagenic double-stranded DNA breaks (DSBs) ^28^ confirmed by a local increase of BLISS (Breaks Labelling In Situ and Sequencing) ^29^ signals at origins. The formation of DSBs at origins is further supported by the colocalization of the DNA end binding factor XRCC5/Ku80 ^30^ with origins (**Fig. 2g**). Interestingly, BRCA1, which physically and functionally interacts with Ku80 to promote classical non-homologous end joining (NHEJ) ^31^ is also enriched at origins, but the key homologous recombination factor RAD51 is depleted (**Fig. 2g**). The preferential use of mutagenic end-joining pathways at these breaks is further supported by origin-associated ID B signature, which is characterised by deletions with microhomology consistent with alternative NHEJ (**Extended Data Fig. 2d**), which is known to act redundantly with classical NHEJ ^32^.

At the onset of the S-phase, the MCM2-7 helicases, loaded at origins during the G1 phase, are activated and the two replicative CMG (Cdc45-Mcm2-7-GINS) helicase complexes translocate in opposite directions. This creates positive torsional constraints in front of the newly activated replication forks, requiring topoisomerase I to ensure continued progression ^33^. Abortive topoisomerase I activity would lead to the accumulation of potentially mutagenic single-stranded DNA breaks (SSBs) in sequences flanking the origins. We found that TOP1 colocalises with the SSB-sensing factor PARP1 at the boundaries of the origins (**Fig. 2h**) but both factors are depleted at the origins themselves. It has been demonstrated that, in yeast, after priming of DNA replication by polymerase α, polymerase δ initiates both leading and lagging strand synthesis and thus performs the bulk of DNA synthesis in the vicinity of the origin before handing over to polymerase ε for ongoing leading strand synthesis ^34–36^. We thus assessed the exposure of constitutive origins to mutational signatures associated with the activity of POLD1 and POLE, which are revealed when the exonuclease domains of these polymerases are disrupted ^37^. We found that the pattern of substitutions is consistent with polymerase δ being the main replicative polymerase active in the first 2 kb around human origins (**Extended Data Fig. 3d**). Further, the accumulation of S33 phosphorylated RPA2 over the background RPA signal in the flanks of the origins suggests local ATR activation indicative of poorly processive and uncoupled DNA synthesis ^38,39^ (**Fig. 2h**). Together this may lead to an increased burden of mismatches and explain the MMR signatures detected at the origin flanks (**Fig. 2e**). The intensity of all analysed ChIP-seq signals at origins correlates with origin efficiency (**Extended Data Fig. 3e**) supporting that DNA repair deficiency and the accumulation of mutations at origins is due to origin activation. Hence the discrete replication initiation sites imprinted at constitutive origins represent hotspots for mutagenesis that have the potential to impact genome evolution and expression.

### Evolution of origin sequences

To investigate the consequences of mutagenesis at constitutive origins on the evolution of the human genome, we devised an *in silico* model in which a library of DNA sequences, calibrated on non-coding human genomic sequences, was evolved according to rules defined by the observed mutation prevalence, pattern and replication strand bias at constitutive origins (**Fig. 3a**). We computed probability density functions (**Extended Data Fig. 4a-c**) that describe the probability that a nucleotide will be mutated based on its position relative to the origin and the nature of that mutation based on the trinucleotide context and exposure to the known SBS signatures, corrected for base composition. We found that our model recapitulates general features of origins observed in the human genome such as local increase of GC content (**Fig. 3b**), base composition skews (**Fig. 3c** and **Extended Data Fig. 4d**) and the presence of origin *cis-*regulatory elements such as CpG islands ^40^ and G-quadruplexes ^41^ (**Extended Data Fig. 4e-f**). This observation supports the hypothesis that the mutagenic processes operational at origins drives origin sequence evolution towards that observed in the genome of modern humans.

**Figure 3.**
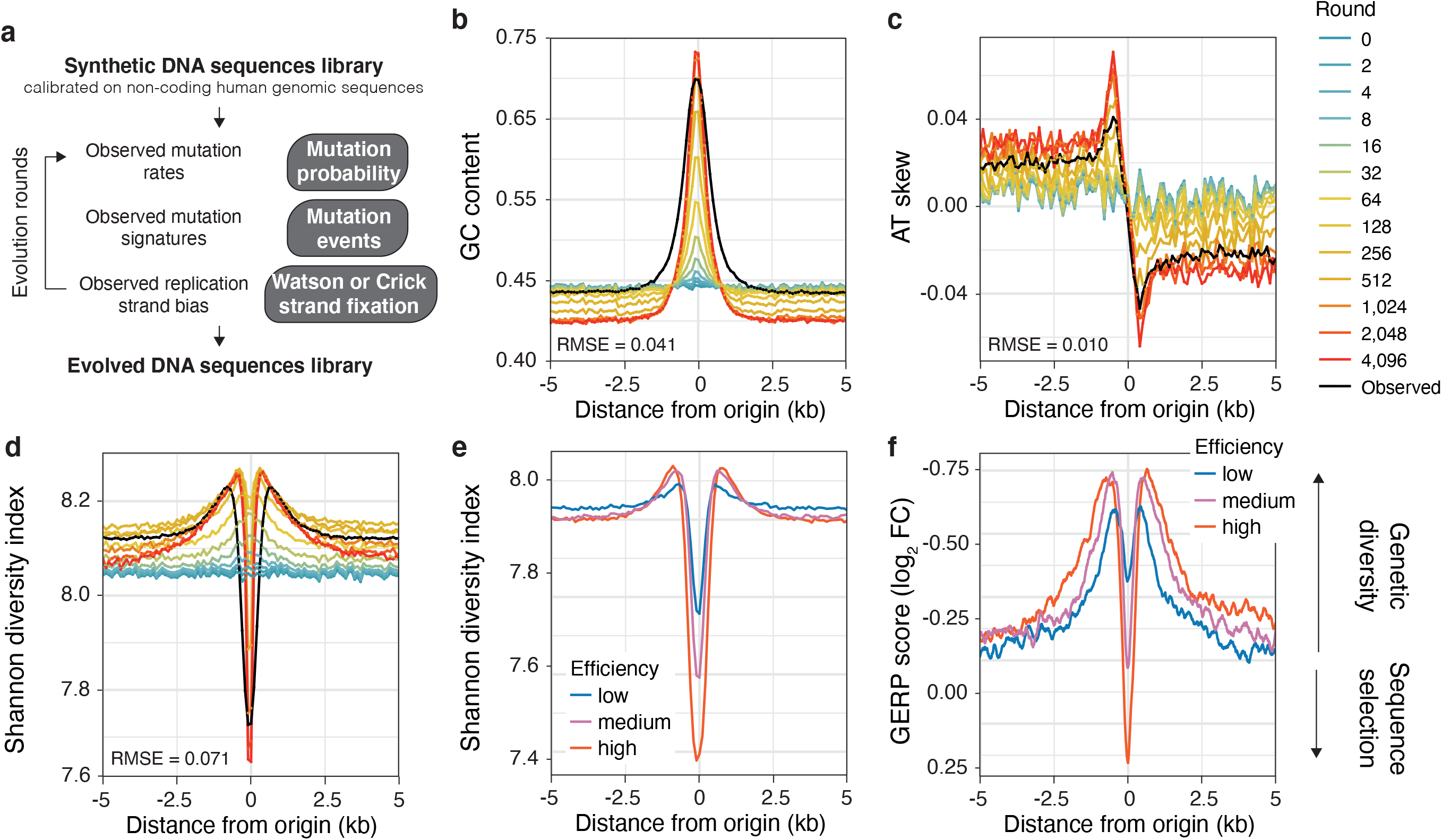
Mutagenesis at origins drives origin specification and genetic diversity. **a**, An *in silico* model was devised to assess the impact of mutagenesis during origin activation on the evolution of sequences at constitutive origins. The model uses rules, see **Methods**, defined on observed mutation rates, mutagenic processes and replication asymmetry at constitutive origins to evolve a library of 1,000 DNA sequences calibrated on non-coding regions of the human genome. Applying sequential rounds of evolution recapitulates features of constitutive origins such as (**b**) local increase in GC content, (**c**) AT skew and (**d**) sequence diversity computed using Shannon diversity indexes. Coloured lines represent values at different rounds of evolution and black lines correspond to values observed at constitutive origins. The Root Mean Square Error (RMSE) values are the standard deviations of the residuals between computed values at round 4,096 and observed values. **e**, Shannon diversity indexes decrease, indicating sequence selection, and increase, indicating sequence diversity, at origin centres and flanks respectively. This is replication dependent as the signal depends on the efficiency of constitutive origins. **f**, Rates of genome evolution, plotted as GERP scores, at constitutive origins are replication-dependent, and display positive and negative values at the centres and flanks of origins respectively. GERP scores were normalised using background values from domains adjacent to origins. Positive and negative GERP scores indicate an increase and decrease in nucleotide substitution rates relative to a genome-wide expectation of neutral evolution respectively. For panel **e** and **f**, origins were binned according to low (blue), medium (purple) or high (red) efficiency.

We next analysed variation in the Shannon diversity index, a measure of sequence diversity, around our modelled *in silico* origins (**Fig. 3d**). Sequence diversity decreases over time at origins but increases in their flanks. The resulting ‘volcano-shaped’ patterns around origins, reminiscent of the exposure distribution of signatures SBS B and C (**Fig. 2b**), suggests that the mutational processes linked to replication initiation have the capacity not only to create the sequence environment observed at replication origins but also to drive genetic diversity in the immediate vicinity of origins. These impacts are dependent on origin efficiency (*P* = 8.26 × 10^−4^, chi-square test of independence, **Fig. 3e**) suggesting that the most efficient sites of replication initiation contribute to the evolution of the human genome. To test this hypothesis, we analysed the rate of genome evolution around constitutive origins by assessing local variation of the genomic evolutionary rate profiling (GERP) score, which quantifies nucleotide substitution rates relative to a genome-wide expectation of neutral evolution ^42^. At origin centres, we found positive GERP scores, indicating purifying selection for functional elements, while in the flanks we observed negative GERP scores, indicating increased nucleotide substitution rates (**Fig. 3f**). Furthermore, increases in substitution rate correlated with origin efficiency (*P* < 2.2 × 10^−16^, chi-square test of independence, **Fig. 3f**) and usage (**Extended Data Fig. 4g**) demonstrating that genetic diversity is linked to the efficiency of replication initiation.

### Origins drive phenotypic diversity

To assess the functional consequences of the increased mutation rate at origins, we analysed the positional enrichment of origins within the human genome (**Fig. 4a**). We found that all origin classes are enriched within genes (*P* < 2.2 × 10^−16^ for all origin classes, chi-square test) with ~ 80 % of constitutive origins lying within gene bodies. Constitutive origins locally accumulate at element boundaries such as transcription start sites (TSS) and splice junctions (**Extended Data Fig. 5a**) suggesting that variants associated with constitutive origins are more likely to have a functional impact. To assess whether this is the case, we used RegulomeDB ^43^ and found a significant enrichment at constitutive origins of SNPs with predicted effects on gene regulation, hereafter refer to as functional variants (RegulomeDB score ≤ 2; a 4.68-fold increase when compared to all SNPs, **Fig. 4b**). This enrichment reflects replication initiation as it increases with origin usage (**Fig. 4b**) and correlates with constitutive origin efficiency (*P* = 7.49 × 10^−206^, chi-square test of independence, **Extended Data Fig. 5b**). We then investigated whether the presence of a constitutive origin within a gene feature increases the mutational load in the associated gene. We first categorised genes according to the presence or absence of a constitutive origin at their TSS or at splicing sites, and computed the density of functional variants around their TSS (**Fig. 4c**) and splice sites (**Fig. 4d**). We found that gene features marked by an origin display increased mutation load compared with those that do not. Thus, genes with an origin within 2.5 kb of their TSS display a ~ 1.55-fold increase in mutational load (**Fig. 4c** and **Extended Data Fig. 5c**) compared with those without, and this enrichment correlates with origin efficiency (*P* = 2.72 × 10^−6^, chi-square test of independence, **Extended Data Fig. 5d**). Similarly splice sites marked by an origin display a ~ 1.34-fold increase in mutational load (**Fig. 4d** and **Extended Data Fig. 5c**) that also corelates with origin efficiency (*P* = 8.3 × 10^−101^, chi-square test of independence, **Extended Data Fig. 5d**). This increase in functional variants at TSS and splice sites marked by constitutive origins suggest that mutagenesis at origins has the potential to be translated into gene expression variation and to modulate mRNA splicing.

**Figure 4.**
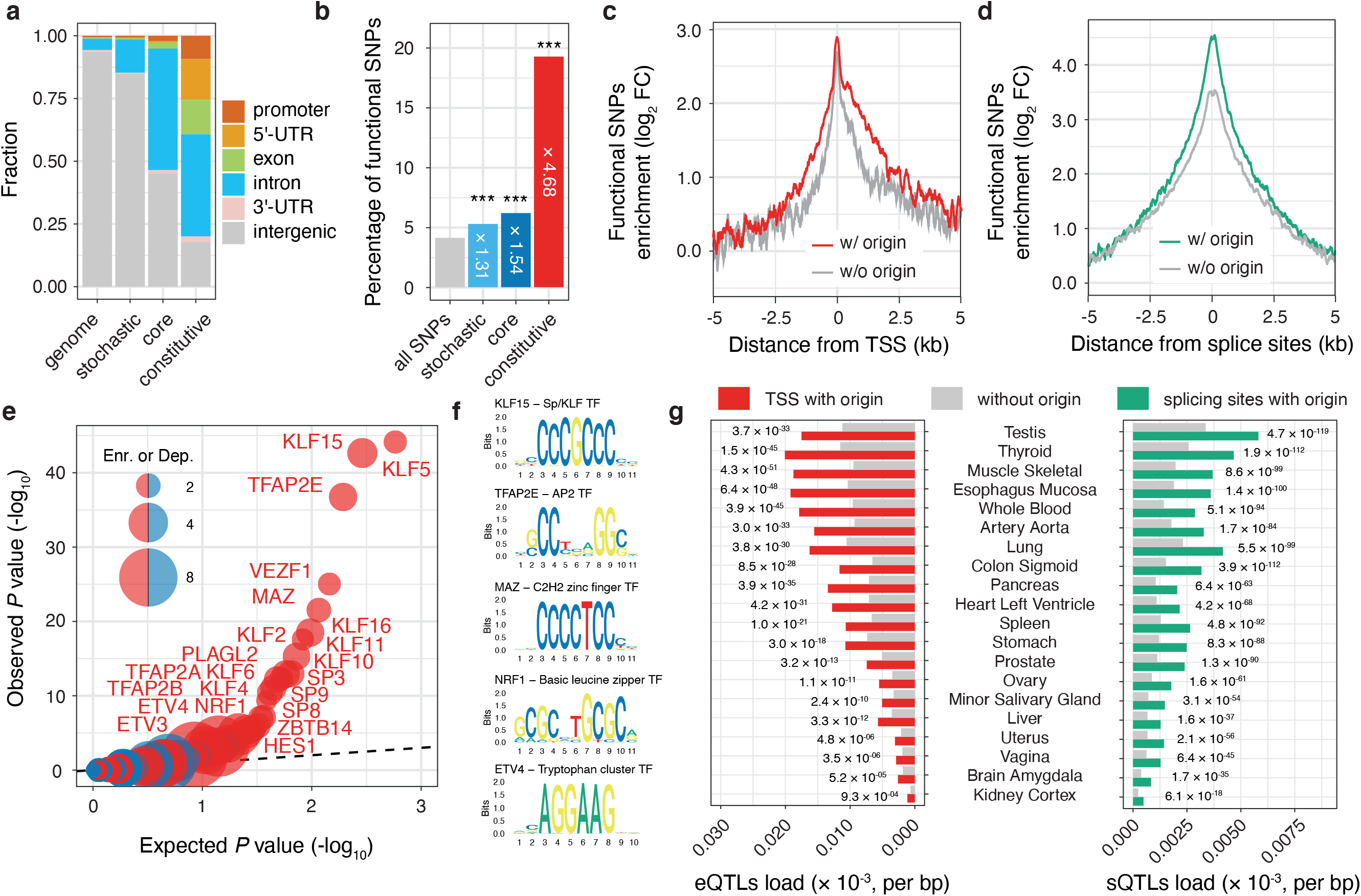
Mutagenesis at constitutive origins drive gene expression variation. **a**, All origin classes are enriched within genes suggesting that variants associated with origins are likely to have a functional impact. The barplot reports the distribution of stochastic, core and constitutive origins within gene features considering origin midpoints. **b-d**, Origin SNPs are predicted to affect gene regulation. **b**, Percentages of variants, found within 5 kb domains centred on origins, predicted to have regulatory potential. Variants with predicted effects on gene regulation were identified using RegulomeDB that leverages signals generated by the ENCODE and other projects to identify functional SNPs according to their overlap with regulatory regions of the human genome. The presence of constitutive origins increases mutational loads associated with functional variants at gene TSS (**c**) and splice sites (**d**). **c**, Protein coding genes were categorized according to the presence (*n* = 11,410) or absence (*n* = 7,925) of constitutive origins within 2.5 kb around their TSS. **d**, Splice sites of GENCODE basic transcripts were categorized according to the presence (*n* = 55,903) or absence (*n* = 121,511) of constitutive origins within 2.5 kb. Mutation densities were computed in bins of 100 bp and normalised using background values from adjacent domains. Mutagenesis at constitutive origins disrupts specific TF binding. **e**, Quantile-quantile plot of origin variants associations with TF binding sites. Enrichment and depletion are reported by red and blue bubbles respectively, which size reflects values. **f**, Functional SNPs at constitutive origins overlap with specific GC-rich TF binding sites. *cis-*eQTLs and *cis-*sQTLs are enriched at the TSS and splice sites of genes, respectively, marked by constitutive origins compared to sites free of origins. **g**, *cis-*eQTLs and *cis-*sQTLs densities within 5 kb regions centred on gene features were computed for features marked by constitutive origins (red and green bars for TSS and splice sites respectively) or free from origins (grey bars) in 49 human tissues. The figure reports a selection of tissues (the full figures are reported in **Extended Data Fig. 6a and 6b**). *P* values are computed using chi-square tests (panel **b, e** and **g**), *** *P* < 2.2 × 10^−16^.

To test whether this is the case, we first asked whether origin-associated SNPs have the potential to alter transcription factor (TF) binding sites. We extracted the sequence context of all functional variants found within 2.5 kb around TSS and created lists of predicted TF binding sites associated with the functional variants. By comparing the enrichment of TF motifs around TSS marked by an origin compared with those that are origin free (**Fig. 4e**) we observed enrichment of GC-rich binding sites (**Fig. 4f**), including those of members of the Sp/KLF and AP-2 families that play a vital role in regulating the growth and development of a large number of tissues ^44,45^. Mutational load at these TF binding sites correlates with origin efficiency (**Extended Data Fig. 5e**) suggesting that mutagenesis at constitutive origins moulds TF binding in a replication-dependent manner. We then investigated whether alteration of TF binding by origin mutagenesis induces gene expression variation in human populations. Using datasets generated by the Genotype-Tissue Expression (GTEx) project ^46^ to identify *cis-*expression quantitative trait loci (*cis-*eQTL) / gene pairs, we analysed mutational loads in 2.5 kb regions around TSS marked by constitutive origins, and those without, and computed *cis*-eQTL enrichment in both sets. For the 49 screened tissues, we found an increase in *cis*-eQTL loads, ranging from 1.70 to 2.87-fold (*P* < 2.7 × 10^−3^, chi-square test, **Fig. 4g** and **Extended Data Fig. 6a**), in gene TSSs marked by an origin. A similar analysis revealed that splice sites marked by an origin display increased *cis*-splicing quantitative trait loci (*cis*-sQTL) loads (up to 2.15-fold with *P* < 6.1 × 10^−18^, chi-square test, **Fig. 4g** and **Extended Data Fig. 6b**) supporting the notion that mutagenesis at constitutive origins also affects mRNA splicing. Taken together these results suggest that genetic variants induced by mutagenesis at constitutive origins are a source of gene expression variation and phenotypic diversity in human populations.

## Discussion

Here we establish a class of highly efficient and mutagenic replication origins that allow the observation of specific mutational patterns associated with replication initiation that substantially impact genome evolution and expression. A direct relationship between origin activation and mutagenesis is supported by the significant correlations between origin usage/efficiency and mutation frequency. We hypothesise that the two mutational processes we identify as operating at constitutive replication origins may result from the two-step process of origin activation (**Fig. 2f**), providing a framework for future experimental validation. Our model suggests that replication initiation would trigger mutagenesis at all initiation sites, but we note that stochastic and core origins exhibit a similar mutation rate than their flanking regions (**Fig. 1c**). This could be due to the lower burden of origin mutagenesis at these sites that is not separable from the background, reflecting that only the most recurrent sites of replication initiation leave a significant mutational imprint.

We demonstrated that mutagenesis at constitutive origins is the driving force for origin sequence evolution, creating local nucleotide skews (**Fig. 3c** and **Extended Data Fig. 4d**) known as S-jumps. S-jumps are found at bacterial, viral and eukaryotic origins ^47,48^, suggesting that the role of origins in shaping genomes is conserved throughout evolution. We note that while constitutive origins cover only ~ 0.7 % of the human genome, their enrichment within gene bodies leads to a high local mutation density with disproportionate functional consequence. From an evolutionary perspective, it may seem paradoxical that a replication initiation-coupled mechanism focusses mutation to promoters or splice sites. However, such targeted mutagenesis may provide a significant advantage in shaping the evolution of the multiple promoters ^49^ and alternative splicing sites ^50^ found in the majority of human genes and thus influence the diversity of tissue-specific mRNA isoforms available for selection of new functions.

Although the mutagenesis we report here reflects evolutionary events occurring during normal gametogenesis and early development, we anticipate that dysregulation of mechanisms that maintain genome stability in cancer will modulate mutagenesis at replication initiation sites and exert an additional mutational burden on genetic elements marked by constitutive origins. Knowing that oncogene induction also activates origins within highly transcribed genes ^51^, we anticipate that mutagenesis arising from origin activity may contribute extensively to the evolution of cancer genomes.

## METHODS

### Human reference genome and annotation

All analyses were performed on the hg38/GRCh38 human reference genome assembly. Data sets originally obtained with coordinates on other assemblies were projected onto the hg38 assembly using ‘liftOver’ from the rtracklayer R/Bioconductor package ^52^ with the corresponding chain files obtained from http://hgdownload.cse.ucsc.edu in the individual assembly download sections. Gene structure annotations were extracted using the biomaRt R/Bioconductor package ^53^. We considered only transcription start sites (TSSs) of protein coding genes (*n* = 19,335) and splice sites of GENCODE basic transcripts (*n* = 59,563) to prioritise full-length coding transcripts over partial or non-coding transcripts. Intron coordinates were recovered by subtracting the coordinates of exons, 5’-UTRs and 3’-UTRs from transcript coordinates using the ‘subtract’ command of bedtools ^54^.

### Replication origins and timing

Replication origins, mapped using a Short Nascent Strand isolation protocol coupled with next-generation sequencing (SNS-seq), from 19 human cell samples were recovered from ref ^14^. Early replications origins, mapped by initiation site sequencing (ini-seq 2), were recovered from reference ^15^. To categorised constitutive, core and stochastic origins, we consider that two initiation sites represent the same origin if the mapped domains are found within less than 5kb. Distances between origins were computed using the ‘closest’ function of bedtools. To demonstrate the effect of origin efficiency on mutagenesis, we defined three groups of constitutive origins, referred to as Low-, Medium- and High-efficiency constitutive origins, by dividing the distribution of observed efficiencies into three quantiles and assigning each origin to one of the three groups. Replication timing was computed using data from ref ^55^ considering the log2 ratio of early- (G1+S1) to late- (S4+G2) replicating DNA values as a function of genomic positions in 50 kb sliding windows at 1 kb intervals. We averaged the obtained timing values from experiments using the GM12812, BG02, IMR-90, MCF7 cell lines. Enrichment of replication origins within replication timing domains was assessed by computing the fraction of origins and associated replication timing values in 100 kb windows of the human genome.

### Mutation data

Whole-genome common population variants were obtained from dbSNP build 151 on hg38. The common subset reports all variants representing alleles observed in the germline with a minor allele frequency of at least 1% and mapped to a single location in the reference genome assembly. Taken as a set, common variants are less likely to be associated with severe genetic diseases due to the effects of natural selection, following the view that deleterious variants are not likely to become common in the population. Single nucleotide polymorphisms (SNPs, *n* = 34,082,281) and short insertions/deletions (indels, *n* = 3,673,496) were recovered using VCFtools ^56^ and analysed independently.

### Mutation rate estimation

In order to compare mutation rates at origins to their neighbouring regions, we considered 10 kb flanks either side of origin mid-points. As the presence of replication origin clusters may bias the mutation rate analyses, we considered only domains that contain a single origin. Analyses were then performed considering domains covering 9,351 constitutive, 12,224 core and 10,923 stochastic origins. Local mutation rates were computed in 100 bp windows covering origin domains as the ratio of the total number of mutations, SNPs or indels, over the total number of considered domains. We corrected the local mutation rates by the basal levels of mutation and considered the increases of mutation rate over background by normalising the previous values by the average values obtained for the first and last 20 windows of origin domains. Mutation interdistances were computed for each mutation as the smallest distance between the first upstream or downstream mutation and we averaged values over 100 bp windows covering origin domains. To assess whether mutation rates at origins are due to the local sequence context, we considered the total number of the six pyrimidine mutations, divided by the number of C/G or A/T for C and T mutations respectively, and considered the enrichment of mutations over background by normalising the previous values by the average values obtained for the first and last 20 windows of origin domains. This corrects for mononucleotide compositional biases that are abundant in the vicinity of replication initiation sites.

### Replication strand bias

The total number of mutations, *n*, corresponding to a given base pair change *b1:b2* → *m1:m2* and its complementary mutation *b2:b1* → *m2:m1* were computed in 100 bp windows covering the 20 kb origin domains previously defined. Replication strand biases, *RSB*, for each window were then calculated as, 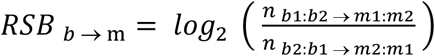. Trends in replication strand biases and inversion of asymmetry at constitutive origins are highlighted using local Loess regression fits.

### Mutation signatures analysis

Evaluation and visualisation of mutational patterns at origins were performed using the MutationalPatterns ^18^ and NMF ^57^ R/Bioconductor packages.

*De novo* extraction of mutational signatures was performed using an unsupervised non-negative matrix factorization (NMF) approach. We defined mutation count matrixes by considering the frequencies of SNPs, calculated in each of the possible 96 trinucleotide 5′ to 3′ contexts, or indels, calculated considering size, nucleotides affected and presence of repetitive and/or microhomology regions, in 100 bp windows spanning the 20 kb constitutive origin domains defined previously. We determined the number of single-base substitution (SBS) and indel (ID) signatures by considering the smallest rank at which the cophenetic correlation coefficient starts decreasing. We thus extracted three SBS and ID mutational signatures using 1,000 iterations to achieve stability and avoid bad local minima. Signature exposures to constitutive, core and stochastic origin domains, but also domains covering Low-, Medium- and High-efficiency constitutive origins, were constructed by considering the relative contribution of the SBS A-C and ID A-C signatures in 100 bp windows spanning the 20 kb origin domains. To assess the replication-dependency of SBS A-C, we considered the relative contribution of each signature to domains around origins where the signatures operate. We considered 1 kb domains, 4 kb domains but omitting the previous domains and 16 kb domains excluding the previous ones centred at the origins for SBS B, C and A respectively.

To determine the underlying molecular mechanisms associated with the extracted SBS A-C signatures, we analysed the contribution of known signatures of somatic mutations in noncancerous and cancerous human tissues, collected by the Catalogue Of Somatic Mutations in Cancer (COSMIC v3.2) and available from https://cancer.sanger.ac.uk/signatures/sbs, to mutagenesis at constitutive origins. In order to identify the minimum set of mutational processes operating at constitutive origins and avoid signature misattribution, we used an iterative fitting approach with the ‘fit_to_signatures_strict’ function of the MutationalPatterns package and a ‘cutoff max_delta’ of 0.004. We then assigned to each operating signature one of the origin-associated signature SBS A-C based on their cosine similarity and exposure. Known signature exposures to constitutive, core and stochastic origin domains, but also domains covering Low-, Medium- and High-efficiency constitutive origins, were constructed by considering the absolute number of mutations attributed to the operating signatures in 100 bp windows spanning the 20 kb origin domains.

To determine the contribution of polymerase ε and δ to DNA synthesis in the vicinity of constitutive origins, we analysed the contribution of mutational signatures attributed to defective proofreading due to acquired mutations in the exonuclease domains of POLE or POLD1. To do this, we used an iterative fitting approach, as described previously but using only the polymerase-associated signatures SBS10a, SBS10b, SBS10c and SBS10d (COSMIC v3.2) and found that SBS10b (defective POLE proofreading) and SBS10c (defective POLD1 proofreading) contributes to mutagenesis at constitutive origins. Calculating the absolute number of mutations attributed to both signatures in 100 bp windows spanning the 20 kb origin domains allows the demonstration that polymerase δ, but not polymerase ε, is active in a small 4 kb domain centred at the origins.

### DNA repair factor enrichment analysis

The distribution of DNA repair factors and double-stranded DNA breaks (DSB) around constitutive origins was assessed using available ChiP-seq and other datasets. The references and associated Gene Expression Omnibus (GEO) accession numbers for the datasets used in these analyses are reported in **Supplementary Table 1**. We selected datasets assessing the chromatin distribution of ORC (ORC1 and ORC2 subunits, *n* = 2), TOP2B (*n* = 2), DSBs (Breaks Labelling In Situ and Sequencing (BLISS), *n* = 3), BRCA1 (*n* = 4), XRCC5 (*n* = 2), RAD51 (*n* = 1), TOP1 (*n* = 3), PARP1 (*n* = 3), RPA (RPA1 and RPA2 subunits, *n* = 2) and phospho-RPA2-S33 (*n* = 1). We then computed coverage plots using the ‘map’ function of bedtools in windows of 100 bp covering 20 kb domains centred on constitutive origins, but also domains covering Low-, Medium- and High-efficiency constitutive origins. Enrichments over background were computed by normalising the local enrichment values by the average values obtained for the first and last 20 windows of origin domains. We finally averaged the coverage plots over the different analysed cell lines and conditions when available.

### *In silico* model of evolution

To determine the effect of mutagenesis at constitutive origins on their sequences, we devised an *in silico* model of evolution in which a library of synthetic DNA sequences was evolved according to rules defined by observed mutation processes operating at constitutive origins. We defined three probability density functions (PDFs) that describe (i) the probability of a base to mutate according to its distance from origin centres (PDF-1, **Extended Data Fig. 4a**), (ii) the probability of the resulting mutation according to its trinucleotide context and the nature of the known mutational signatures operating at its position (PDF-2, **Extended Data Fig. 4b**) and (iii) the probability for a mutation to be fixed on the top or bottom strand (PDF-3, **Extended Data Fig. 4c**). The library of 20 kb DNA sequences contained 1,000 sequences calibrated on non-coding upstream sequences from the hg38 version of the human genome and using a Markov model with an oligonucleotide size of 6 (http://rsat.sb-roscoff.fr/random-seq_form.cgi). PDF-1 was constructed by fitting a multi-peak gaussian profile to the observed mutation rates reported in **Fig. 1b**. PDF-2 was defined by fitting the contribution of known signatures of somatic mutations in noncancerous and cancerous human tissues (COSMIC v3.2) using a SNP mutation count matrix, describing mutagenesis at constitutive origins, corrected by mononucleotide and trinucleotide composition. This corrects for the excess of base substitutions that are due to compositional biases and allows modelling the impact of replication initiation on ‘naïve’ sequences. Mutational signature exposures were then fitted using a combination of gaussian or sigmoidal functions according to the observed profiles. PDF-3 was constructed by fitting a combination of gaussian or sigmoidal profiles to the observed replication strand biases reported in **Fig. 1d**. We then apply 5,000 rounds of evolution to the DNA library by using PDF-1 to select 100 bases to mutate and PDF-2 and PDF-3 to output the set of mutated sequences. The *in silico* evolution process was stopped when values associated with general features of constitutive origins, such as the GC content, plateaued.

### Sequence features

DNA sequence analysis was performed using the seqinr ^58^ and Biostrings ^59^ R/Bioconductor packages. Nucleotide composition, nucleotide skews and diversity indexes were computed in 100 bp windows spanning 20 kb centred on origin mid-points. AT and GC skews were computed as: 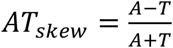 *and* 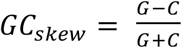, where A, T, C and G are the number of the corresponding nucleotides in the considered window. G-quadruplexes were predicted by considering the G_3+_N_1– 7_G_3+_N_1–7_G_3+_N_1–7_G_3+_, where N refers to any bases, regular expression ^60^. Shannon (or Shannon–Wiener) diversity indexes (SDI) were computed by considering the occurrence of each of the k-mers (k = 6) and the following equation: 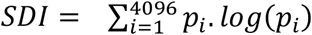, where *p*_*i*_ is the proportional abundance of one of the 4,096 6-mers. As SDI values depend on the number of sequences analysed, computed values have been corrected to account for different sample size when appropriate.

### Human conservation measures

Genomic evolutionary rate profiling (GERP) scores ^42^ were used as a measure of nucleotide diversity between species. The single-nucleotide resolution bigWig file (gerp_conservation_scores.homo_sapiens.GRCh38.bw), reporting substitution rates computed from the multiple alignment of 111 mammalian genomes, was obtained from ftp://ftp.ensembl.org/pub/release101/compara/conservation_scores/111_mammals.gerp_conservation_score/. GERP scores were computed as averages from rolling windows of 100 nt spanning 20 kb centred on origin mid-points. For consistency of presentation with plots of mutation rates and sequence diversity, the y axes in plots showing GERP scores have been been inverted so that greater constraint is low and greater diversity is high.

### Annotation of functional variants

To test whether origin-associated SNPs are likely to have a functional impact on gene expression, we used RegulomeDB ^43^. RegulomeDB annotates SNPs with known and predicted regulatory elements within the human genome. Known and predicted regulatory DNA elements include regions of DNase hypersensitivity, binding sites of transcription factors, and promoter regions that have been biochemically characterized to regulate transcription. Sources of these data include public datasets from GEO, the ENCODE project, and published literature. Rank scores for common SNPs (regulomedb_dbsnp153_common_snv.tsv) were obtained from https://regulomedb.org/regulome-help/. A RegulomeDB rank ≤ 2 was used to predict SNPs with the minimal functional evidence. This resulted in the identification of 524,563 functional SNPs out of the 13,275,023 referenced SNPs.

### Transcription factor binding sites enrichment

To assess the ability of constitutive origin-associated SNPs to disrupt binding of transcription factors (TFs), we extracted the sequence context (SNP ± 12 nt) of all functional variants, as previously defined, found within 5 kb of TSSs and identified overlapping TF binding sites (TFBSs) using the TFBSTools ^61^ R/Bioconductor packages. We used the JASPAR 2020 database ^62^ to curate a list of 639 non-redundant human transcription factors with associated Position Weight Matrixes (PWMs). We then analysed each functional SNP sequence context for TFBS using the ‘searchSeq’ function of TFBSTools using a minimum score of 95% and no strand information. TFBSs were considered as overlapping with a functional SNP if the match scores exceeded an arbitrary value of 10 and the consensus sequences define by the PWMs contain a functional SNP. Enrichment of TFBSs at TSSs marked by origin was then tested by comparing the TFBS/SNP pair lists associated with TSS marked or free of origins. *P*-values associated with the enrichment or depletion of TFBSs were computed using 2 × 2 contingency chi-square tests and compared to the normal expected uniform distribution to identify TFBS disrupted by constitutive origin-associated functional variants. Replication-dependent mutational loads at TFBS were then assessed by computing the number of functional variants overlapping with TFBS hits at TSSs marked by low-, medium- and high-efficient constitutive origins.

### Quantitative trait loci analysis

To characterise the impact of mutagenesis at constitutive origins on human phenotypic diversity, we analysed mutational loads for molecular *cis*-quantitative trait loci (*cis*-QTL) at TSS and splice sites marked by constitutive origins. Expression quantitative trait loci (eQTL, GTEx_Analaysis_v8_eQTL.tar) and splicing quantitative trait loci (sQTL, GTEx_Analaysis_v8_eQTL.tar) for 49 tissues were recovered from the Genotype-Tissue Expression (GTEx) project website at https://www.gtexportal.org/home/datasets. QTL/gene pairs were defined by identifying significant variant-gene associations with *qval* ≥ 0.05 and *cis*-QTL as variants found within 5 kb domains centred at TSSs or splice sites. *cis*-QTL loads were computed as the ratio of the number of significant *cis-*QTLs over the number of bases covered by the considered domains, *i*.*e*. TSS and splice sites marked or free of constitutive origins. *P*-values associated with the enrichment of *cis-*QTLs at TSS and splice junctions marked by constitutive origins were computed using 2 × 2 contingency chi-square tests.

### Computational and statistical analyses

Analysis and all statistical calculations were performed in R (version 4.0.3).

## Code availability

The R codes used to analyse the data and generate the figure reported in the manuscript are available for download at https://github.com/Sale-lab/OriMut

## Acknowledgments

The authors thank M. Babu, J. Yeeles, T. Krude and the members of the Sale group for the comments on the manuscript. Work in the Sale group is supported by a core grant to the LMB by the MRC (U105178808).

## Author contributions

P. M. designed the project and performed the experiments. All authors analysed and interpreted the data. P. M. and J. E. S. wrote the manuscript.

## Competing interests

The authors declare no competing interests.

## EXTENDED DATA FIGURES LEGENDS

**Extended Data Fig. 1.**
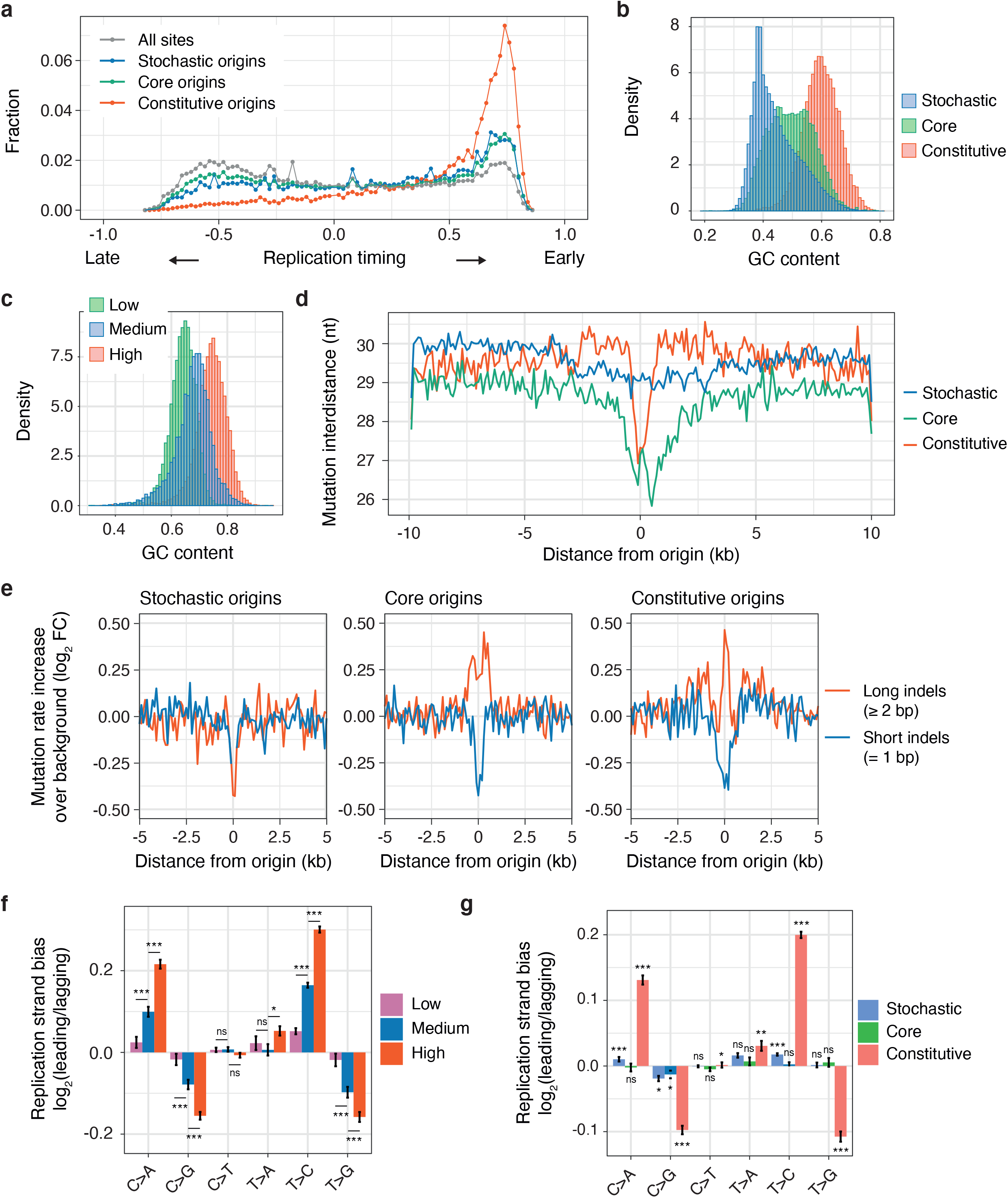
Constitutive origins are hotspots for replication-dependent mutagenesis. **a**, Human replication origins were categorised as stochastic and core origins according to their human cell types specificity. A subset of core origins, mapped by initiation site sequencing (ini-seq 2), were found to be enriched within early replicating domains of the human genome. Origin base composition determines origin usage (**b**) and efficiency (**c**). Origins considered in panel **c** are subsets of constitutive origins from panel **b** binned in three groups of equal sizes and different efficiencies. The density plots, reporting the GC content of 1 kb or 100 bp, for panel **b** and **c** respectively, sequences centred on origins, show that constitutive origins display higher GC contents than core and constitutive origins and that highly-efficient origins are more GC rich that the less efficient ones. **d**, Local decrease in SNP interdistances at origins compared with their flanking domains (*P* = 1.68 × 10^−8^, 7.79 × 10^−24^ and 1.15 × 10^−5^ for constitutive, core and stochastic origins respectively, chi-square test). Mutation interdistances were computed for each mutation as the smallest distance between the first upstream or downstream mutation and we averaged values for all variants in 100 bp windows covering origin domains. **e**, Mutation rates for long (≥ 2 bp) but not short (= 1bp) indels is increased at core and constitutive origins compared to their flanking domains (*P* = 2.82 × 10^−5^ and 3.73 × 10^−4^ for constitutive and core origins respectively, chi-square test). Mutation rates are reported as the increase over background values computed from domains adjacent to origins. Replication strand biases for variants at origins are efficiency- (**f**) and usage- (**g**) dependent. Replication strand biases were computed, as the ratio between the density of mutation at a given base pair over the density of mutation at the complementary base pair in 100 bp windows covering 20 kb origin domains as in **Fig. 1d**. We then computed the ratio between averaged values for 10 kb domains covering leading- (left of origins) over lagging- (right of origins) strand synthesis. *P* values for the comparison of the distribution of individual values computed in 100 bp windows were calculated using the Kolmogorov-Smirnov test, *n*.*s*. non-significant, \**P* < 0.05, \*\**P* < 0.01 and \*\*\**P* < 0.001.

**Extended Data Fig. 2.**
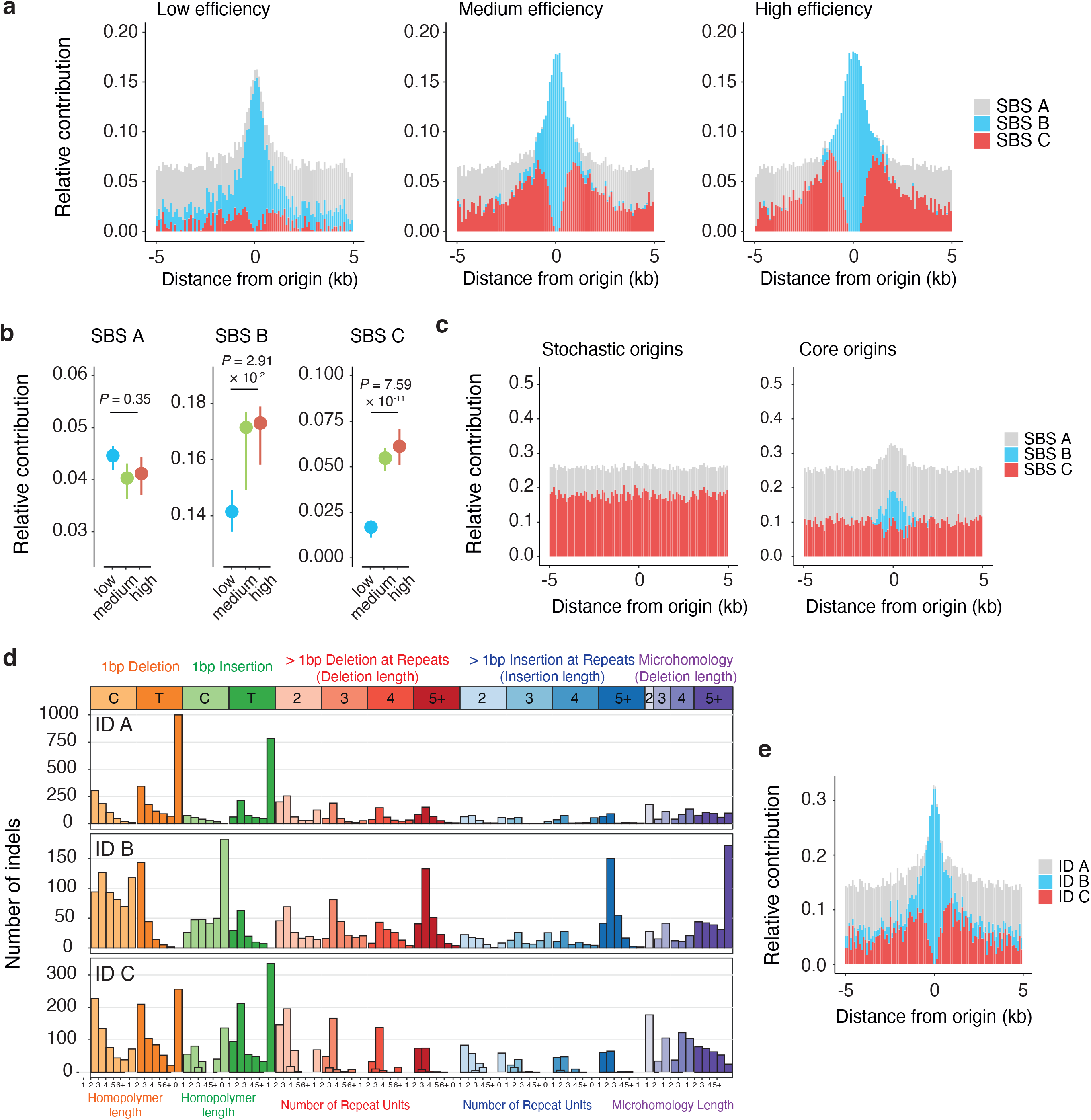
Replication-dependent mutational processes operating at constitutive origins. **a**, The contribution of the *de novo* extracted SBS B and C signatures to mutagenesis at constitutive origins correlates with origin efficiency. Reporting the relative contribution of SBS A-C at constitutive origins binned as low-, medium- and high-efficient origins shows that while the contribution of SBS A remain constant over origin domains, the absolute contribution of SBS B and C increases with origin efficiency. **b**, To assess the replication-dependency of SBS A-C, we considered the relative contribution of each signature to domains around origins where the signatures operate. We considered 1 kb domains, 4 kb domains but omitting the previous domains and 16 kb domains excluding the previous ones centred at the origins for SBS B, C and A respectively. Reported *P* values are from chi-square tests of independence assessing the impact of origin efficiency on SBS A-C contribution. **c**, The contribution of SBS B and C to mutagenesis at origins reflects origin usage with any contribution to mutagenesis at stochastic and core origins largely swamped by background signal from SBS A. **d**, An unbiased *de novo* mutational signature analysis identified three small insertion/deletion (indel) signatures, ID A-C, operating at constitutive origins. Each profile reports the number of mutations attributed to each indicated indels type. ID A and C signatures are characterised by insertions and deletions at long (≥ 5 bp) homopolymers. ID B signature is characterised by increased long (≥ 5 bp) deletions with at least 5 bp of microhomology at their boundaries. **e**, Computing the relative contribution of ID A-C to mutagenesis at constitutive origins shows that ID A-C operate in concert with SBS A-C as they present similar profile (see **Fig. 2b**).

**Extended Data Fig. 3.**
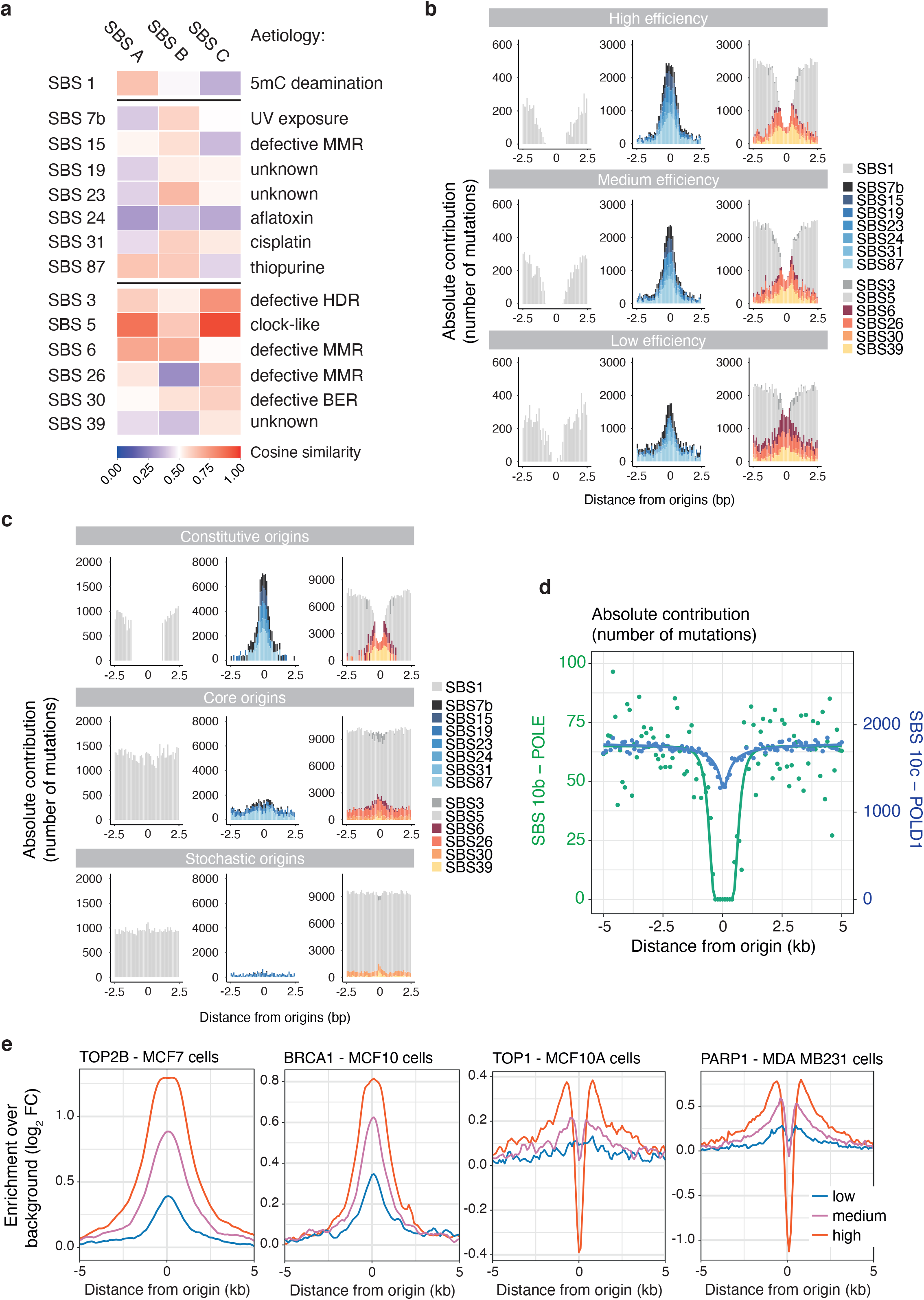
Characterisation of the molecular mechanisms underlying mutagenesis at constitutive origins. Known signatures of somatic mutations in noncancerous and cancerous human tissues, collected by the Catalogue Of Somatic Mutations in Cancer (COSMIC v3.2), were fitted to mutational count matrixes computed from the frequencies of SNPs observed at constitutive origins. We used an iterative fitting approach to select the minimum set of mutational processes operating at constitutive origins and avoid signature misattribution. We identified a set of 14 known signatures that mostly reconstruct the mutation profiles of SBS A-C. **a**, Comparison of the *de novo* extracted SBS A-C signatures to the set of 14 known signatures based on cosine similarity. These signatures were assigned to one of the SBS A-C signature according to their similarity and exposure. Exposure of origins to the mutational processes defined by these SBS signatures is replication-dependent as their contribution to origin mutagenesis is efficiency- (**b**) and usage- (**c**) dependent. We plotted the absolute contribution of the 14 known signatures split in three groups to highlight the three mutational processes described by SBS A-C (left, middle and right columns show the contribution of SBS signatures reconstructing the profiles of SBS A, B and C respectively). **d**, Absolute contribution of signature SBS 10b and SBS 10c, associated with the activity of POLE and POLD1 respectively. Both signatures are attributed to defective proofreading due to acquired mutations in the exonuclease domains of the polymerases. The pattern of substitutions is consistent with polymerase δ being the main replicative polymerase active in the first 2 kb around origins. The number of mutations attributed to both signatures were computed in 100 bp windows spanning the 20 kb origin domains and profiles were fitted using sigmoidal functions. **e**, Replication-dependent enrichment of TOP2B, BRCA1, TOP1 and PARP1 at constitutive origins. Coverage plots were computed from available ChIP-seq datasets (**Supplementary Table S1**) in windows of 100 bp covering 20 kb domains centred on constitutive origins categorised as Low-, Medium- and High-efficient origins. Enrichments over background were computed by normalising the local enrichment values by the average values obtained for the first and last 20 windows of origin domains.

**Extended Data Fig. 4.**
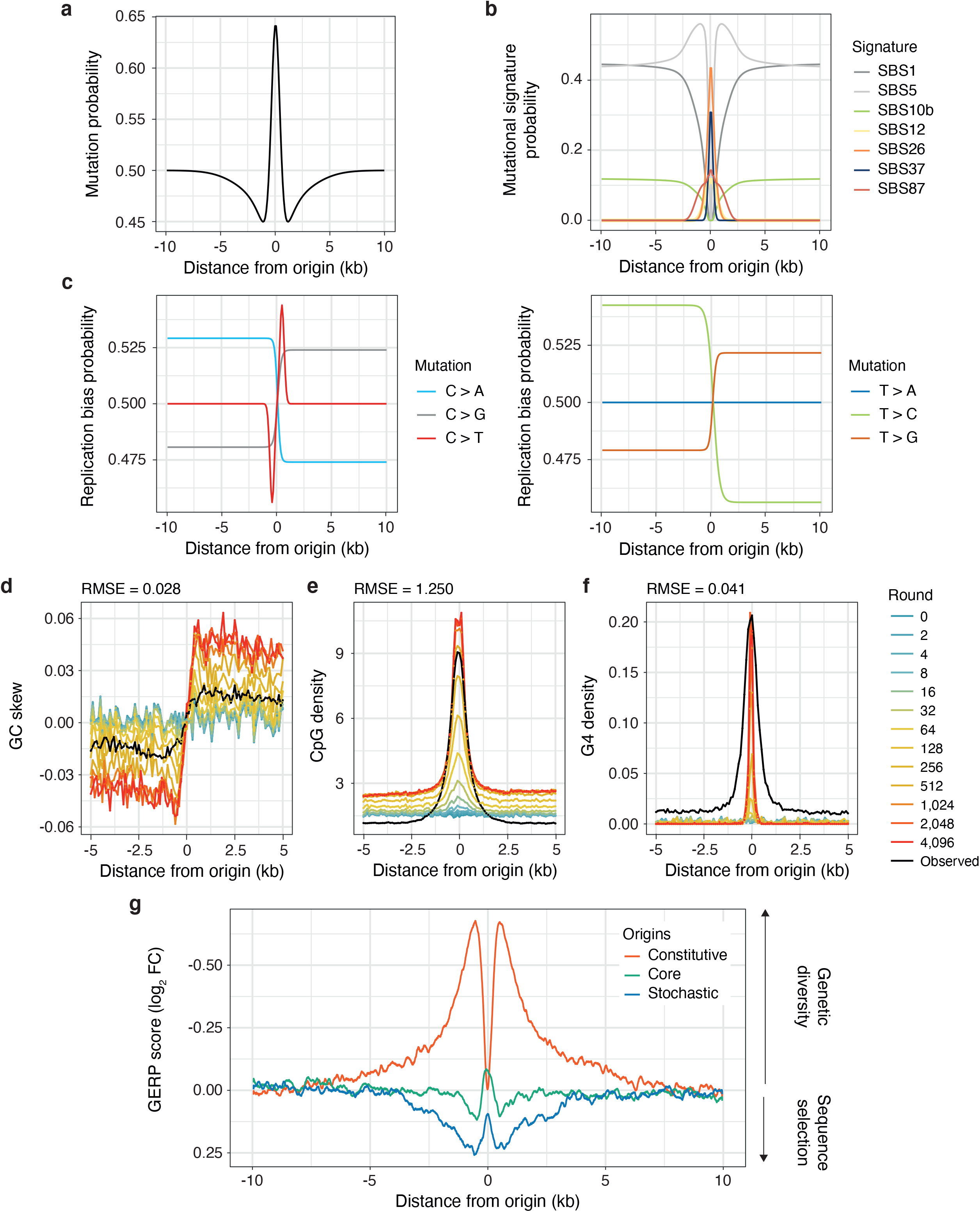
An *in-silico* model of evolution recapitulates general features of constitutive origins. To determine the effect of mutagenesis at constitutive origins on origin sequences, we devised an *in silico* model of evolution in which a library of synthetic DNA sequences was evolved according to rules defined by observed mutation processes operating at constitutive origins. We defined three probability density functions (PDFs) that describe (**a**) the probability of a base to mutate according to its distance from the origin centre, (**b**) the probability of the resulting mutation according to its trinucleotide context and the nature of the known mutational signatures operating at its position and (**c**) the probability for a mutation to be fixed on the top or bottom strand (See **Methods** section for details). A library of 1,000 × 20 kb DNA sequences calibrated on non-coding upstream sequences from the human genome were evolved using these PDFs to select the bases to mutate and output the resulting mutated sequences. 5,000 rounds of *in-silico* evolution allow recapitulation of the features of constitutive origins such as (**d**) GC-skews, (**e**) CpG and (**f**) G-quadruplexes (G4s) density. Coloured lines represent values at different rounds of evolution and black lines correspond to values observed at constitutive origins. The Root Mean Square Error (RMSE) values are the standard deviations of the residuals between computed values at round 4,096 and observed values. **g**, Rates of genome evolution, plotted as GERP scores, at constitutive, core and stochastic origins. GERP scores were normalised using background values from domains adjacent to origins. Positive and negative GERP scores indicate an increase and decrease in nucleotide substitution rates relative to a genome-wide expectation of neutral evolution respectively.

**Extended Data Fig. 5.**
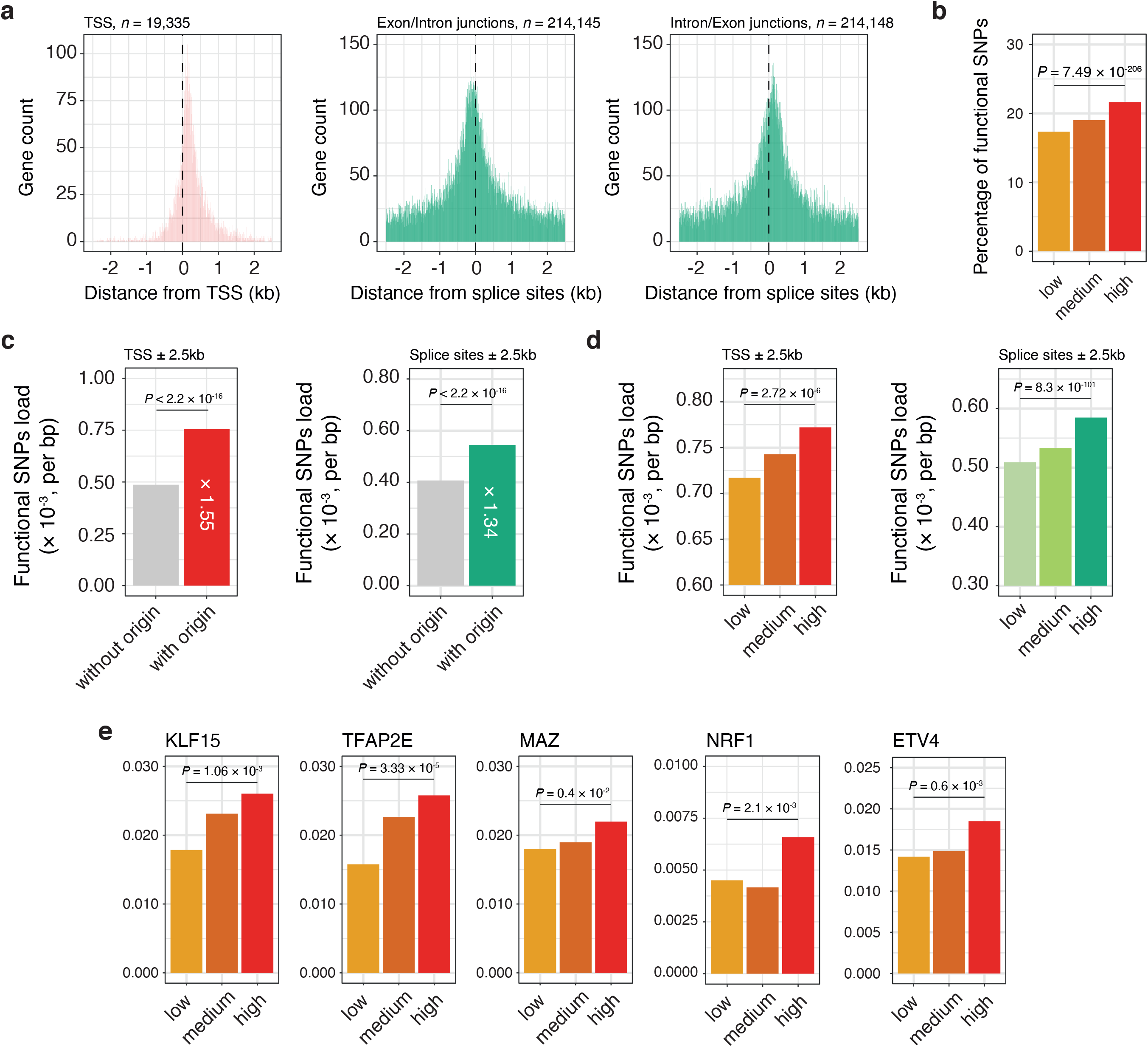
Mutagenesis at constitutive origins drive gene expression variation in humans. **a**, While constitutive origins are enriched within gene bodies (see **Fig. 4a**), we did not find enrichments in specific gene features but rather an accumulation at element junctions such as TSSs, Exon/Intron and Intron/Exon junctions suggesting an impact of mutagenesis at origins on gene expression and mRNA splicing. We considered only protein coding genes (*n* = 19,335) and GENCODE basic transcripts (*n* = 59,563) to prioritise full-length coding transcripts over partial or non-coding transcripts. **b**, The percentage of functional SNPs associated with constitutive origins is replication-dependent as it correlates with origin efficiency. **c**, The presence of constitutive origins increases mutational loads associated with functional variants at gene TSS and splice sites. Quantification of functional SNPs load was performed considering 5 kb domains centred on either TSS or splice sites. **d**, The percentage of functional SNPs associated with constitutive origins at TSS and splice sites is replication-dependent as it correlates with origin efficiency. **e**, Mutational loads at transcription factor (TF) binding sites associated with constitutive origin functional variants correlates with origin efficiency. Selected TFs are enriched at gene TSSs marked by constitutive origins when compared with those that are origin free (see **Fig. 4e**). *P* values were calculated using chi-square tests of independence.

**Extended Data Fig. 6.**
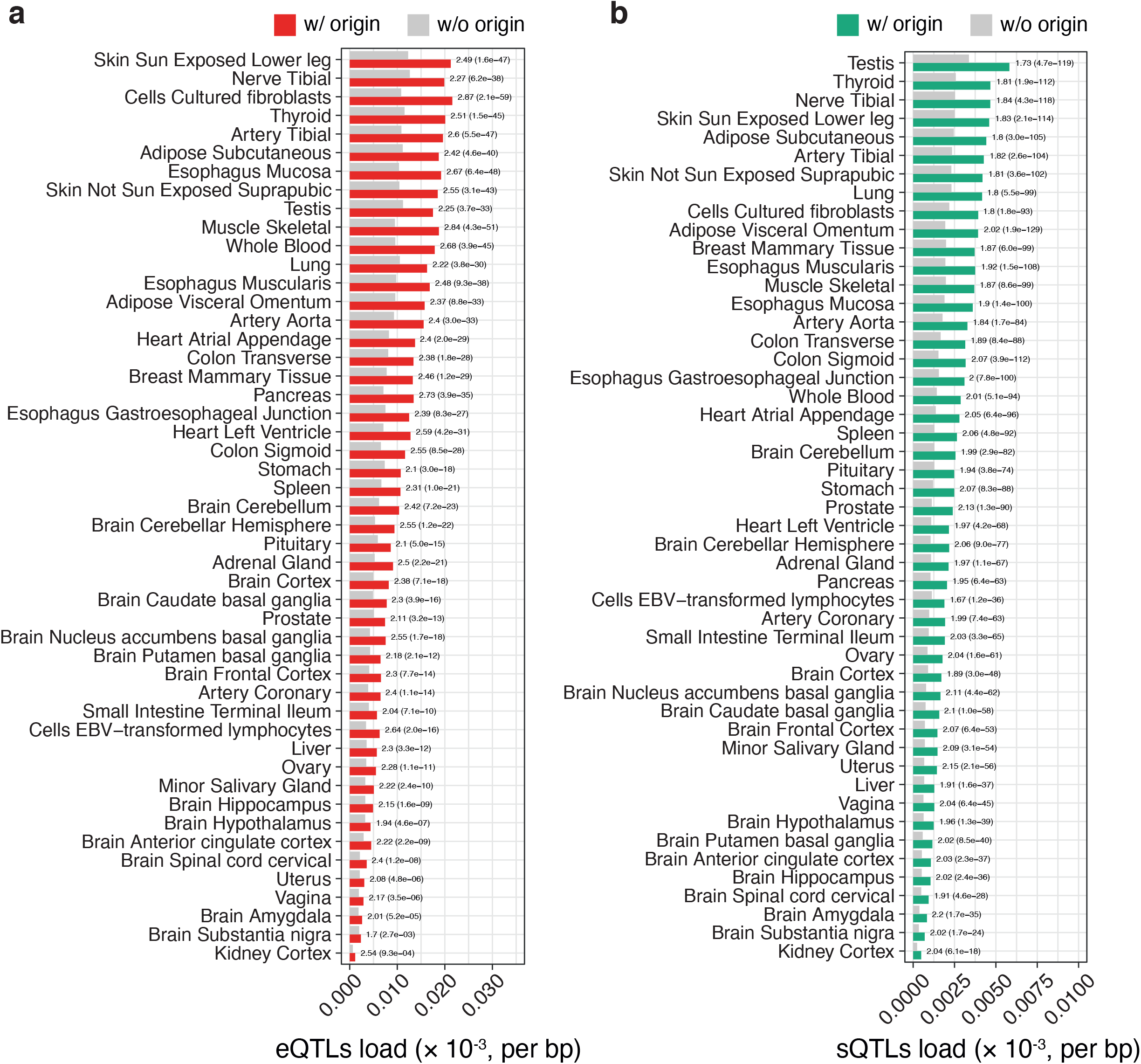
Mutagenesis at constitutive origins drive gene expression variation and alternative splicing in humans. **a**, *cis-*expression quantitative trait loci (*cis-*eQTL) and (**b**) *cis-*splicing quantitative trait loci (*cis-*sQTL) densities within 5 kb regions centred on gene TSS or splice sites when computed for features marked by constitutive origins (red and green bars for TSS and splice sites respectively) or free from origins (grey bars) in 49 human tissues together with enrichment values. This panel reports the full analysis related to **Fig. 4g**. *P* values were computed using chi-square tests.

**Supplementary Table 1.** Gene Expression Omnibus (GEO) accession numbers and associated references for the datasets used in the analyses reported in **Fig. 2g-h** and **Extended Fig. 3e**.

| Target | Cell line | GEO accession numbers | GSM accession numbers | References |
| --- | --- | --- | --- | --- |
| ORC1 | HeLa S3 | GSE37583 | GSM922790 | 63 |
| ORC2 | K562 | GSE70165 | GSM1717888 | 64 |
| TOP2B | MCF7 | GSE141528 | GSM4205700 | 65 |
| TOP2B | MCF10 | GSE93038 | GSM2442946 | 66 |
| BLISS | TK6 | GSE121740 | GSM3444984 | 67 |
| BLISS | K562 | GSE121740 | GSM3444988 | 67 |
| BLISS | CD34 | GSE121740 | GSM3687236 | 67 |
| BRCA1 | MCF10A | GSE40591 | GSM1340576 | 68 |
| BRCA1 | H1 | GSE31477 | GSM935517 | ENCODE Project |
| BRCA1 | HeLa | GSE51334 | GSM935552 | ENCODE Project |
| BRCA1 | GM12878 | GSE31477 | GSM935377 | ENCODE Project |
| XRCC5 | K562 | GSE120110 | GSM3393608 | ENCODE Project |
| XRCC5 | HepG2 | GSM3391789 | GSM3393560 | 69 |
| RAD51 | MCF10A | GSE93038 | GSM2442936 | 66 |
| TOP1 | MCF10A | GSE93038 | GSM3182665 | 66 |
| TOP1 | HCT116 | GSE57628 | GSM2058666 | 70 |
| TOP-seq | HCT116 | GSE57628 | GSM1385717 | 70 |
| PARP1 | MCF10 | GSE93038 | GSM3182666 | 66 |
| PARP1 | MCF7 | GSE61916 | GSM1517305 | 71 |
| PARP1 | MDA-MB231 | GSE61916 | GSM1517306 | 71 |
| RPA1 | HeLa | GSE76661 | GSM2033106 | 72 |
| RPA2 | HeLa | GSE76661 | GSM2033104 | 72 |
| phospho-RPA2-S33 | HeLa | GSE108172 | GSM2891676 | 73 |

## Notes

### Competing Interest Statement

The authors have declared no competing interest.

